## Appendix. Supplementary materials for methods and discussion for "Novel methodology for assessing total recovery time in response to unexpected perturbations while walking"

#### Section 1: Methods

##### 1.1. Steady state walking velocity estimation

The VR controller was programmed to define the steady state walking velocity attained after the following: (1) minimum 30 s of walking, and (2) 12 s duration during which there was a walking speed coefficient of variance < 2%. Steady state walking velocity is determined by a real-time algorithm that is fed treadmill speed data estimated directly from an encoder on the drive shaft of the treadmill motor, which provides velocity signals from the running belts (< 0.1% error).

##### 1.2. Experimental procedures related to the larger protocol – not reported in the present manuscript

Prior to the main experiment in the VR system, pen and paper cognitive tasks (color trails and digit span tests) were administered to assess sustained attention and divided attention [1], along with the NeuroTrax (NeuroTrax Corporation, NY, USA) computerized battery of cognitive tests [2] which assesses different cognitive domains related to motor and postural control. Fall history was also recorded. Participants learned and practiced a serial number addition task, which was the basis for the dual task cognitive load condition. They were instructed to listen to list of 15 numbers that were read to them at a pace of one per two seconds (based on the Paced Auditory Serial Addition Task [3]) and to report the sum of the numbers after the reading. An over ground walking test (10 meter walk test, [4]) and time up and go test [5] were conducted along with a high level functional test (HiMAT: High Level Mobility Assessment Tool,

[6,7]). Electroencephalography (EEG), electromyography (EMG), electrocardiography (ECG), respiration, and galvanic skin response (GSR) sensors were placed on participants, who were also fitted with a safety harness and 41 reflective markers before stepping onto the treadmill (see Fig. 1, main text).

#### 1.3. Paradigm

The following conditions were implemented. (1) Postural stability (P1, P2, P3, P4): P1 – quiet standing with eyes open (60 sec), P2 – quiet standing with eyes closed (30 sec), P3 – quiet standing with eyes open and performing the serial number addition task (30 sec), P4 – quiet standing with eyes closed and performing the serial number addition task (30 sec); (2) Gait task (G1, G2, G3): G1 – participant taught how to walk in self-paced mode, at preferred speed; G2 – walking in self-paced mode with perturbations (single task condition, 5 min, performed three times), G3 – walking in self-paced mode with perturbations while performing the serial number addition task (dual task condition, 5 min, performed three times).

In condition G1, participants were instructed to start walking and report when they had reached their preferred speed. After achieving this, participants were asked to dynamically change their gait speed, including coming to a full stop and resuming walking. This was repeated until the participants felt comfortable with the process and until the examiner, a trained physical therapist (UR), felt that the participants were performing the transitions from standing to preferred walking speed smoothly. Conditions G2-G3 were implemented in random order among participants. Walking conditions were implemented in two different VR scenes: a park and a suspended bridge.

#### 1.4. Detailed explanation of the operation of the third stage of the total recovery time detection algorithm

Since our aim was to identify the window where 'no further significant reduction in amplitude is detected',i.e. Eq. S-1 is fulfilled (see Methods, main text), an iterative process was executed during the third stage of operation of the total recovery time algorithm. The process included scanning a moving window (i.e., 20 values of the OSDev starting at perturbation onset), computing the 'amplitude' within each window, and 'deciding' based on Eq. S-1 if the '$best_{ind}$' (index of the first point in the window containing the best amplitude difference, $best_{amp}$, see below) had been detected. We continued this process until the moving window contained the last data point, to check whether the $best_{ind}$ required updating (i.e., if there was a window containing a 'new' best amplitude). At the heart of this process are the criteria shown in Eq. S-1, which set the condition if the current amplitude ($cur_{amp}$) and current index ($cur_{ind}$) are to be assigned to $best_{amp}$ and $best_{ind}$, respectively. In other words, the algorithm scans each window (called the 'current window' in each iteration) starting at perturbation onset and moves to the right, and if an amplitude is found that is satisfying Eq. S-1, the algorithm sets the current amplitude as the best amplitude.

(Eq. S-1)

$$\frac{current\ gain\ from\ improvement}{gain\ from\ first\ amp\ improvement} = \frac{best_{amp} - cur_{amp}}{first_{amp} - best_{amp}} > 1\% \cdot (cur_{ind} - best_{ind})$$

where $first_{amp}$ = the amplitude of the first window right after perturbation (the a priori assumption is that this is the largest amplitude difference due to the reaction to perturbation); $cur_{amp}$ = the amplitude of the current window; and $best_{amp}$ = an amplitude that is at least 50% smaller than the first amplitude. For the first window, $best_{amp}$ is an arbitrary value. The $best_{amp}$ is reevaluated every iteration for update (eventually the recovery point is the starting position of this window). Thus, the $best_{amp}$ is updated when two conditions are met: (1) a smaller amplitude difference has been found, i.e, $cur_{amp} < best_{amp}$; and (2) the "profit" from an update exceeds the "cost" of delaying the recovery point. This "cost" is based on the index difference of $cur_{amp}$ and $best_{amp}$; practically, we used a threshold of 1% for the "profit versus cost" ratio (see heuristic example in Fig. S1 C).

Intuitively, the decrease from $best_{amp}$ to $cur_{amp}$ should be significant compared to the decrease from $first_{amp}$ to the present $best_{amp}$. Furthermore, the formula in Eq. S1 is stricter if the position of the current window is further away from the $best_{amp}$ window index. This is done in order to overcome the natural variance of amplitudes. The point of recovery is defined as the $best_{ind}$ after calculating all amplitude differences in the whole OSDev vector within consecutive windows of 20 samples. Fig. S1 depicts the process of defining the point of recovery.

Prior to seeking the point of recovery, the algorithm assesses whether there was a detectable effect of the perturbation on the gait parameters of step length and step width. The criterion for 'no deviation' (i.e., no detectable effect) works on the OSDev. We computed five amplitude differences in four windows prior to perturbation and in the first window after perturbation (which always contains the maximum amplitude difference in OSDev due to the perturbation). From the four amplitudes prior to perturbation, we obtained the characteristic amplitude of the a priori behavior of OSDev, i.e., mean and standard deviation of the amplitudes (M_apri_amp and SD_apri_amp, respectively). In the case that the maximal amplitude (i.e., the first amplitude after perturbation) does not exceed M_apri_amp+2*SD_apri_amp, the algorithm will output ‘no deviation’ (i.e., no deviation from previous behavior was detected as a result of the perturbation). Next, the algorithm scans the OSDev to find the point of recovery. The general idea is that as long as the amplitude keeps decreasing significantly, the point of recovery has not yet been reached. When no further significant change in amplitude is detected, the point of recovery is designated (green arrow and circle in Figs. 4A and C, main text, respectively). Total recovery time is then determined via subtraction of perturbation time from point of recovery time.

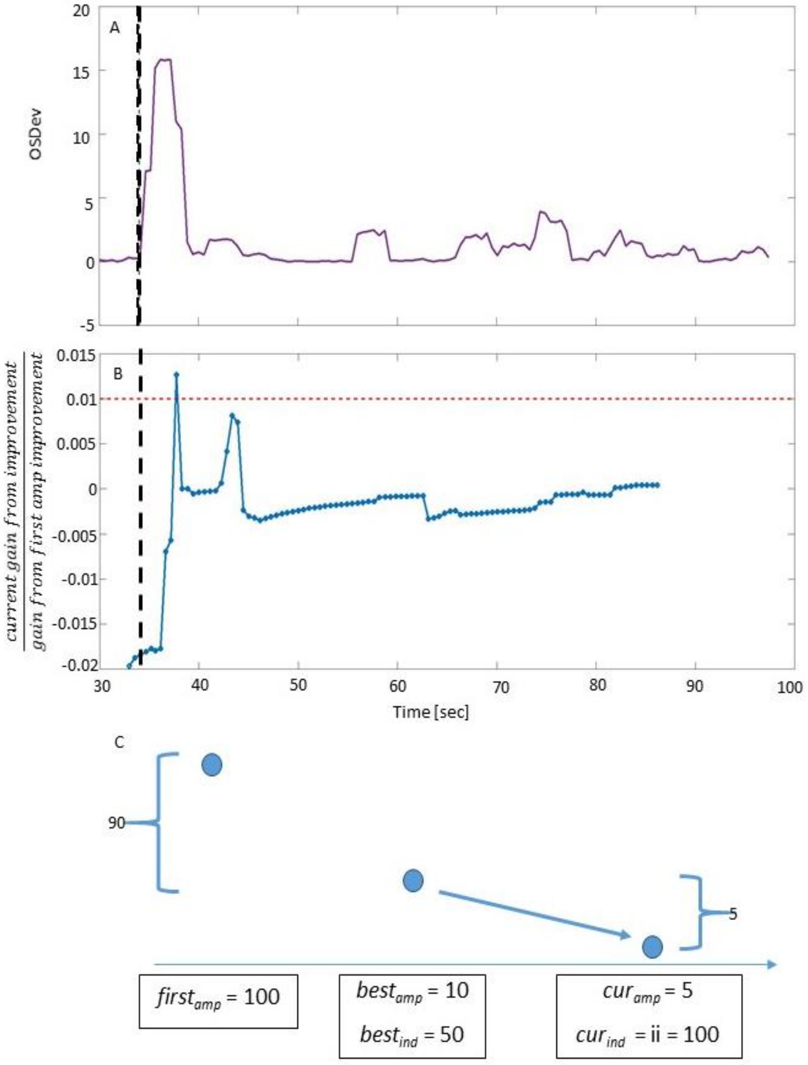

**Fig. S1. Illustration of the algorithm's assessment if $best_{amp}$ should be updated with $cur_{amp}$ (same perturbation as in Fig. 4, main text).**

(A) The overall sum of deviations graph (OSDev; same as panel C of Fig. 4, main text). Black dashed line represents perturbation onset. (B) The ratio $\frac{\textit{current gain from improvement}}{\textit{gain from first amp improvement}}$ (see Eq. S1; blue line) is compared with 1% (red dashed line). If the ratio crosses the red line, $best_{amp}$ is updated. Since no such crossing occurs after data point 8 (at ~ 37 sec), the point of recovery is designated at that point. (C) A heuristic example of the evaluation if the $cur_{amp}$ should be updated to $best_{amp}$ (i.e., one data point example of how we calculate the graph in panel B). If the $first_{amp}$ is 100, the $best_{amp}$=10, $best_{ind}$=50, but current amplitude = 5, $cur_{ind}$ = 100, the algorithm will use Eq. S1 to assess whether or not to update the $best_{amp}$. The first improvement in this case is 90 (100-10). The current amplitude improves to 5 (10-5), which is negligible in comparison to 90 (5/90 → 5.5%), so the algorithm will not update the recovery point.

### Section 2: Results

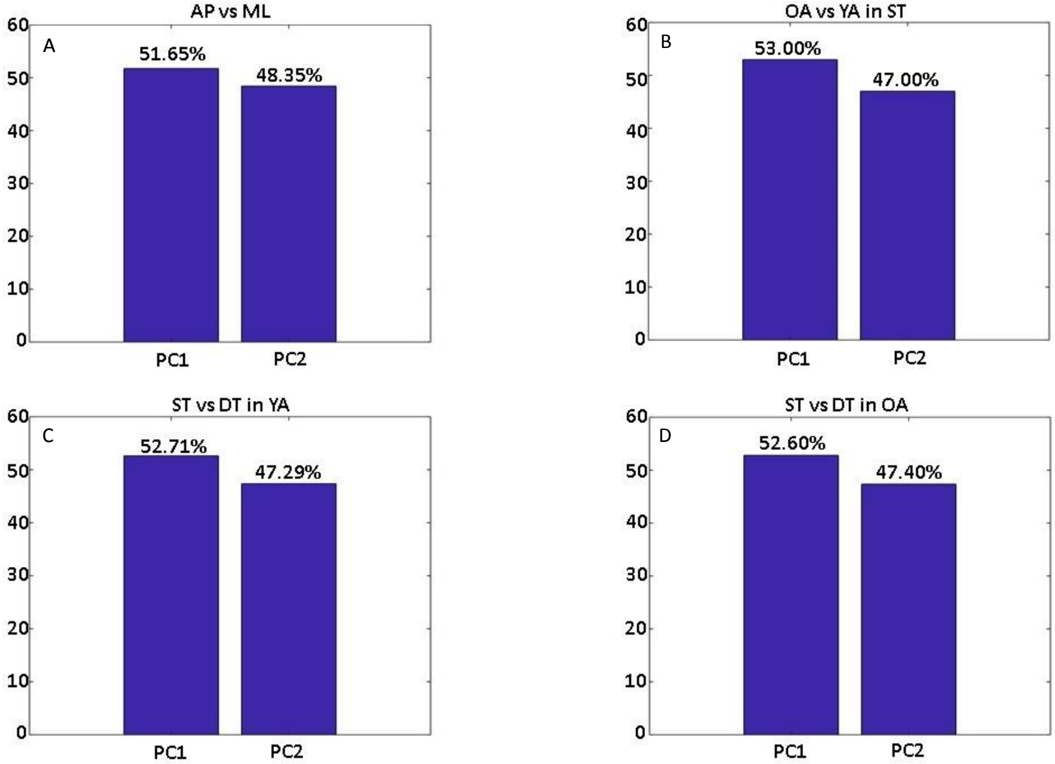


**Fig. S2:** Scree plots representing the principal components and the variance explained by them for total recovery time values of step length and step width for the following comparisons: (A) perturbation direction – anterior-posterior versus medio-lateral; (B) group – older adults versus young adults; (C) **cognitive** load conditions – single task versus dual task among older adults; and (D) cognitive load conditions – single task versus dual task among younger adults. PC1 and PC2 are the two components found by the PCA and the percentages represent the percent of variation explained in the data by each component.

OA = older adults; YA = young adults; AP = anterior-posterior; ML = medio-lateral; ST = single task; DT = dual task; PC = principal component.
