## Appendix.Table S1 for "Novel methodology for assessing total recovery time in response to unexpected perturbations while walking"

**Table S1. Median and range of recovery times, for Level 12 mid-stance perturbations, by group, condition, and perturbation group type.**

| | | Medio-Lateral Perturbations | | | | | | | | Perturbations Antero-Pesterior | | | |
|---|---|---|---|---|---|---|---|---|---|---|---|---|---|
| | | PLtMsLt | | PLtMsRt | | PRtMsLt | | PRtMsRt | | TmMsLt | | TmMsRt | |
| | | SL | SW | SL | SW | SL | SW | SL | SW | SL | SW | SL | SW |
| OA ST | N | 22 | 22 | 26 | 26 | 22 | 22 | 25 | 25 | 8 | 5 | 19 | 17 |
| | median | 5.00 | 5.25 | 4.88 | 5.70 | 4.63 | 5.67 | 5.03 | 6.03 | 4.13 | 5.96 | 4.60 | 7.20 |
| | min | 3.81 | 3.81 | 3.58 | 4.00 | 3.57 | 3.57 | 3.50 | 4.25 | 3.66 | 5.45 | 3.74 | 3.75 |
| | max | 21.13 | 13.99 | 14.47 | 22.07 | 13.42 | 9.85 | 27.61 | 16.93 | 4.71 | 10.61 | 8.31 | 28.18 |
| OA DT | N | 37 | 37 | 21 | 23 | 10 | 10 | 21 | 21 | 22 | 11 | 18 | 15 |
| | median | 4.67 | 5.19 | 5.38 | 6.30 | 4.68 | 5.18 | 5.32 | 5.08 | 4.55 | 5.78 | 4.59 | 7.40 |
| | min | 3.82 | 3.74 | 3.56 | 4.15 | 3.30 | 4.62 | 3.62 | 4.06 | 3.49 | 3.28 | 3.81 | 3.86 |
| | max | 17.72 | 14.25 | 28.55 | 21.13 | 7.61 | 6.08 | 10.97 | 11.19 | 10.19 | 12.39 | 19.93 | 25.77 |
| YA ST | N | 26 | 29 | 18 | 19 | 25 | 26 | 18 | 23 | 21 | 18 | 32 | 27 |
| | median | 5.37 | 5.36 | 5.00 | 7.13 | 4.77 | 5.63 | 5.92 | 5.59 | 4.77 | 7.19 | 4.47 | 5.38 |
| | min | 4.38 | 4.32 | 3.83 | 4.22 | 3.49 | 4.30 | 4.40 | 4.18 | 4.05 | 4.09 | 2.81 | 3.83 |
| | max | 15.56 | 11.17 | 15.25 | 18.32 | 24.80 | 23.49 | 20.89 | 16.22 | 9.42 | 21.18 | 10.20 | 25.30 |
| YA DT | N | 24 | 27 | 20 | 21 | 21 | 20 | 29 | 33 | 29 | 24 | 27 | 18 |
| | median | 5.06 | 5.93 | 4.83 | 7.43 | 4.51 | 5.82 | 5.18 | 5.49 | 4.88 | 5.99 | 4.62 | 5.57 |
| | min | 4.31 | 4.07 | 3.61 | 4.54 | 3.85 | 4.31 | 3.97 | 4.33 | 3.09 | 3.38 | 3.81 | 2.62 |
| | max | 25.94 | 20.58 | 12.72 | 17.93 | 25.51 | 25.01 | 9.27 | 17.85 | 18.86 | 22.73 | 24.47 | 22.40 |

Min = minimum; Max = maximum; OA= older adults; YA = young adults; ST = single task; DT = dual task; PLtIcLt = platform left initial contact left; PLtIcRt = platform left initial contact right; PRtIcLt = platform right initial contact left; PRtIcRt = platform right initial contact right; TmIcLt = treadmill initial contact left; TmIcRt = treadmill initial contact right; PLtToLt = platform left toe off left; PLtToRt = platform left toe off right; PRtToLt = platform right toe off left; PRtToRt = platform right toe off right; TmToLt = treadmill toe off left; TmToRt = treadmill toe off right; SL = step length; SW = step width.
