## Appendix.Table S2 for "Novel methodology for assessing total recovery time in response to unexpected perturbations while walking"

**Table S2. Median and range of recovery times, for Level 12, initial contact and toe off perturbations, by group, condition, and perturbation group type.**

| | | Medio-Lateral Perturbations | | | | | | | | | | | | | | | | Anterior-Posterior Perturbations | | | | | | | |
|---|---|---|---|---|---|---|---|---|---|---|---|---|---|---|---|---|---|---|---|---|---|---|---|---|---|
| | | PLtIcLt | | PLtIcRt | | PLtToLt | | PLToRt | | PRtIcLt | | PRtIcRt | | PRtToLt | | PRtToRt | | TmIcLt | | TmIcRt | | TmToLt | | TmToRt | |
| | | SL | SW | SL | SW | SL | SW | SL | SW | SL | SW | SL | SW | SL | SW | SL | SW | SL | SW | SL | SW | SL | SW | SL | SW |
| OA ST | Median (n) | 5.88 (5) | 4.96 (5) | 5.52 (3) | 5.73 (3) | 4.26 (3) | 9.00 (3) | 5.78 (6) | 5.29 (6) | 7.43 (3) | 15.11 (3) | 9.15 (6) | 11.92 (6) | 5.33 (5) | 5.60 (5) | 10.86 (2) | 5.53 (2) | 4.90 (1) | [] (0) | [] (0) | [] (0) | 4.72 (4) | 6.33 (3) | 5.25 (8) | 5.92 (7) |
| | Min | 5.22 | 3.94 | 4.72 | 5.52 | 3.69 | 7.15 | 4.61 | 4.53 | 6.33 | 4.57 | 6.33 | 4.57 | 4.51 | 4.50 | 6.39 | 4.67 | 4.90 | [] | [] | [] | 3.82 | 5.74 | 3.76 | 2.02 |
| | Max | 9.89 | 11.84 | 13.21 | 9.51 | 5.53 | 12.02 | 10.53 | 11.49 | 10.87 | 17.08 | 12.71 | 17.08 | 14.44 | 14.68 | 15.32 | 6.39 | 4.90 | [] | [] | [] | 10.57 | 11.57 | 11.17 | 12.79 |
| OA DT | Median (n) | 4.92 (5) | 6.41 (5) | [] (0) | [] (0) | 3.94 (2) | 5.14 (2) | 8.08 (7) | 5.51 (7) | 4.42 (3) | 6.51 (3) | 4.97 (7) | 5.94 (7) | 4.89 (7) | 4.89 (7) | 5.80 (5) | 10.13 (5) | 9.86 (2) | 5.57 (1) | 9.78 (1) | 7.02 (1) | 5.38 (3) | 8.72 (3) | 4.80 (7) | 7.36 (7) |
| | Min | 4.18 | 4.63 | [] | [] | 3.37 | 5.11 | 4.50 | 4.36 | 3.68 | 4.42 | 3.62 | 4.36 | 4.58 | 4.10 | 3.50 | 4.50 | 8.27 | 5.57 | 9.78 | 7.02 | 2.34 | 5.17 | 3.46 | 5.75 |

| | | | | | | | | | | | | | | | | | | | | | | | | | |
|---|---|---|---|---|---|---|---|---|---|---|---|---|---|---|---|---|---|---|---|---|---|---|---|---|---|
| | Max | 9.26 | 12.01 | [] | [] | 4.51 | 5.17 | 17.72 | 9.03 | 9.06 | 8.84 | 9.06 | 8.84 | 7.07 | 5.48 | 11.86 | 18.48 | 11.45 | 5.57 | 9.78 | 7.02 | 6.16 | 20.18 | 17.94 | 24.12 |
| YA ST | Median (n) | 5.33 (1) | 5.91 (1) | [] (0) | [] (0) | 5.64 (1) | 5.06 (1) | 6.58 (6) | 5.53 (6) | [] (0) | [] (0) | 4.66 (2) | 7.49 (2) | 5.13 (6) | 5.74 (6) | 4.48 (4) | 10.22 (4) | 3.46 (2) | 22.82 (1) | 5.59 (1) | 5.59 (1) | [] (0) | [] (0) | 18.74 (1) | [] (0) |
| | Min | 5.33 | 5.91 | [] | [] | 5.64 | 5.06 | 4.27 | 3.68 | [] | [] | 3.90 | 5.42 | 4.22 | 4.83 | 4.03 | 4.26 | 3.06 | 22.82 | 5.59 | 5.59 | [] | [] | 18.74 | [] |
| | Max | 5.33 | 5.91 | [] | [] | 5.64 | 5.06 | 6.87 | 7.50 | [] | [] | 5.42 | 9.56 | 10.52 | 6.85 | 5.16 | 21.63 | 3.87 | 22.82 | 5.59 | 5.59 | [] | [] | 18.74 | [] |
| YA DT | Median (n) | 5.52 (1) | 14.55 (1) | 4.65 (1) | 4.65 (1) | 4.71 (4) | 4.45 (4) | 4.97 (5) | 5.53 (5) | 4.30 (3) | 6.50 (3) | 6.40 (4) | 6.81 (4) | 9.70 (6) | 5.21 (5) | 4.64 (7) | 5.75 (7) | 9.75 (2) | 5.99 (3) | 3.63 (2) | 19.20 (1) | 4.32 (1) | 8.08 (1) | 4.89 (1) | 10.31 (1) |
| | Min | 5.52 | 14.55 | 4.65 | 4.65 | 4.41 | 4.41 | 4.63 | 4.32 | 4.20 | 4.70 | 4.20 | 4.70 | 6.73 | 4.57 | 4.18 | 4.18 | 4.52 | 2.40 | 3.54 | 19.20 | 4.32 | 8.08 | 4.892 | 10.31 |
| | Max | 5.52 | 14.55 | 4.65 | 4.65 | 4.99 | 11.12 | 24.27 | 7.25 | 8.51 | 11.57 | 14.64 | 11.57 | 11.95 | 7.82 | 6.35 | 17.39 | 14.98 | 22.72 | 3.72 | 19.20 | 4.32 | 8.08 | 4.892 | 10.31 |

Min = minimum; Max = maximum; OA= older adults; YA = young adults; ST = single task; DT = dual task; PLtIcLt = platform left initial contact left; PLtIcRt = platform left initial contact right; PRtIcLt = platform right initial contact left; PRtIcRt = platform right initial contact right; TmIcLt = treadmill initial contact left; TmIcRt = treadmill initial contact right; PLtToLt = platform left toe off left; PLtToRt = platform left toe off right; PRtToLt = platform right toe off left; PRtToRt = platform right toe off right; TmToLt = treadmill toe off left; TmToRt = treadmill toe off right; SL = step length; SW = step width.
